# Gray Matters: An Efficient Vision Transformer GAN Framework for Predicting Functional Network Connectivity Biomarkers from Brain Structure

**DOI:** 10.1101/2024.01.11.575307

**Authors:** Yuda Bi, Anees Abrol, Sihan Jia, Zening Fu, Vince D. Calhoun

## Abstract

The field of brain connectivity research has undergone revolutionary changes thanks to state-of-the-art advancements in neuroimaging, particularly regarding structural and functional magnetic resonance imaging (MRI). To navigate the intricate neural dynamics, one must possess a keen comprehension of the interdependent links between structure and function. Such relationships are understudied as they are complex and likely nonlinear. To address this, we created a new generative deep learning architecture using a conditional efficient vision transformer generative adversarial network (cEViTGAN) to capture the distinct information in structural and functional MRI of the human brain. Our model generates functional network connectivity (FNC) matrices directly from three-dimensional sMRI data. Two pioneering innovations are central to our approach. First, we use a novel linear embedding method for structural MRI (sMRI) data that retains the 3D spatial detail. This embedding is best for representative learning, and when used on a consistent dataset, and shows that it is good at upstream classification assignments. To estimate neural biomarkers, we need to process much smaller patches using ViT-based architectures, which usually makes the computations more difficult because of the self-attention operations. We present a new, lightweight self-attention mechanism to address this challenge. Our mechanism not only overcomes computational shortcomings of traditional softmax self-attention but also surpasses pure linear self-attention models in accuracy and performance. This optimization enables us to analyze even the tiniest neuroanatomical details with exceptional precision. Our model allows for the identification of functional network connectivity (FNC) with 74.2% accuracy and also predicts subject differences in FNC for schizophrenia patients versus controls. The results are intriguing and suggest the links between gray matter volume and brain function may be stronger than previously considered.

## I. Introduction

The complex relationship between the brain’s structural and functional characteristics is a subject of significant interest, particularly due to its implications for understanding brain health and disorders, ultimately impacting human quality of life. Structural MRI (sMRI) and functional MRI (fMRI) are two prevalent neuroimaging tools, each providing distinct insights into the brain’s physiology in both health and disease [1], [2], [3]. sMRI is instrumental in revealing high-resolution images of the brain, which is crucial for detecting morphological changes associated with neurological conditions. For example, cortical atrophy and hippocampal shrinkage are key indicators in Alzheimer’s disease, detectable through sMRI [4], [5]. On the other hand, fMRI offers a dynamic view of the brain, identifying active regions during specific tasks or states. This functionality is essential for understanding disorders such as schizophrenia or autism, where brain activity patterns are often altered [6].

The interaction between structural changes and functional abnormalities in the brain is intricate and bidirectional. Structural alterations can precipitate functional impairments, impacting cognitive processes and behavior. Conversely, functional adaptations, driven by factors such as neural plasticity and learning experiences, may induce structural changes. Although this interconnection is acknowledged, it remains not fully elucidated due to individual variances and the influence of genetic and environmental factors [7]. The challenge lies in finding methodologies or technologies that can effectively unravel these complex connections.

In this context, the rapid advancement of artificial intelligence, particularly deep learning, offers promising potential in neuroimaging and diagnosis. Data fusion techniques in deep learning have been employed to explore the correlations between structural and functional brain imaging [8]. However, most research in this area has primarily focused on using multimodal information for predicting behavioral outcomes [9] or diagnosing brain disorders [10]. There is a notable gap in studies aiming to use brain structure to predict brain function. Nevertheless, existing evidence suggests a significant correlation between inter-subject variations in brain structural networks and those observed in resting-state fMRI networks [11]. This indicates an underexplored yet potentially fruitful area of research in understanding the complex interplay between the brain’s structure and function. Generative deep learning models, such as generative adversarial networks (GANs) and variational autoencoders (VAEs), have shown significant success in neuroimaging applications. They excel at simulating neuroimaging data, detecting disease-specific patterns, and enabling modality translation, as exemplified by converting T1-weighted to T2-weighted MRI scans. This proficiency suggests the existence of a potential unifying biological or structural principle in brain imaging, bridging the gap between different imaging techniques [12], [13], [14], [15], [16]. Despite their distinct methodologies, these imaging modalities may converge on a shared informational nexus within the brain, advocating for an integrated approach to understanding brain functions. However, the integration and synthesis of sMRI and fMRI data, particularly in deep learning applications, remain comparatively underexplored [17], [18]. This contrasts with the substantial advancements in GAN models that predominantly focus on medical image synthesis between different modalities.

The preference for medical image synthesis across modalities such as CT, MRI, and PET is driven by multiple factors. Primarily, there is a clinical need to merge the unique benefits of each modality: CT’s detailed bone imaging, MRI’s nuanced soft tissue contrast, and PET’s metabolic insights. This integrative approach aims to enhance diagnostic accuracy and optimize treatment planning. On the other hand, the synthesis of sMRI and fMRI images is less prevalent due to several challenges. The key issue stems from the inherent differences in these modalities: sMRI yields high-resolution anatomical details, essential for structural analysis, while fMRI maps brain activity and cerebral blood flow, pertinent to functional studies. This divergence in focus leads to varying image characteristics and data types, complicating the synthesis process. Additionally, the intricate nature of fMRI data necessitates advanced analytical methods to decode dynamic brain activities, posing a further obstacle to its integration with sMRI. Furthermore, as highlighted earlier, sMRI and fMRI typically serve distinct clinical and research purposes. The former is predominantly used for structural evaluation, whereas the latter is oriented towards functional assessments. This divergence in applications diminishes the immediate clinical impetus to synthesize these two data forms.

The emergence of the vision transformer (ViT) marks a significant shift in image processing, particularly within the realm of neuroimaging [19]. This innovative model diverges from traditional convolutional neural networks (CNNs) by incorporating mechanisms initially crafted for language processing, such as the self-attention mechanism, enabling a more nuanced and holistic image analysis [20]. ViT’s architecture, which dissects images into patches for processing through multiple transformer layers, allows for an in-depth analysis, independent of an image segment’s spatial location. However, its computational demands, particularly its *O*(*N* ^2^) complexity, pose challenges, propelling the quest for more streamlined architectures [21]. As researchers explore innovations to bolster ViT’s efficiency, such as architecture pruning and knowledge distillation, without sacrificing performance [22], [23], the model’s potential within neuroimaging becomes increasingly evident. In this domain, the precision required for detailed brain scans necessitates advanced, lightweight models, highlighting the suitability of ViT’s enhanced interpretability through attention mechanisms.

What is particularly striking is ViT’s prowess at outperforming conventional CNNs, especially when applied to intricate medical imaging datasets. This superior performance raises a pivotal question: could ViT effectively replace CNNs in the backbone of GAN models? This substitution could harness the detailed representational power of ViTs, offering a more refined approach, especially beneficial for intricate studies involved in brain science, such as saliency map research. By leveraging the precision and computational efficiency of ViT, there’s potential for groundbreaking advancements in identifying and visualizing specific biomarkers in brain regions, thus bridging the gap between structural and functional neuroimaging insights.

The significance of conducting research using ViT and GAN to determine the relationship between the structure and function of the human brain is illustrated by the preceding two questions. This paper describes 1) the creation of a new conditional GAN model, which is called cEViT-GAN, that can generate functional connectivity matrices from sMRI data. As the generator and discriminator, an efficient ViT model is used. Similar to the previous question, when providing data before the linear embedding layers, we use tiny regions, which may result in more tokens but can increase the precision of attention maps. 2) In contrast to conventional self-attention operations, we select a block-wise self-attention layer that significantly reduces the computational cost without compromising performance. The mechanism for block-wise self-attention is versatile. The model accentuates regional relationships by employing self-attention operations in each block, thereby capturing localized patterns and interactions. In contrast, when inter-block self-attention is enabled, it ensures that long-term dependencies across the larger structure are not neglected. This dual strategy is ideal for identifying brain biomarkers from structural MRI data. Focusing on specific regional relationships that indicate certain conditions or abnormalities is essential, but it is also necessary to analyze the entire brain image to completely comprehend and diagnose the issue. 3) To increase the efficacy of training, we generate the linear embeddings using a previously trained model based on previous classification tasks. This improves the generator’s training efficiency and effectiveness. 4) In addition, our GAN model has the potential to be used as a biomarker identification tool for identifying the structural and functional connections of the human brain, particularly for various brain diseases such as schizophrenia.

## II. Related Works

Generative adversarial networks (GANs), initially proposed by [24] and extended by [25], have significantly advanced as essential AI tools in various generative tasks. These tasks include image and signal generation [26], as well as text-to-image and image-to-image synthesis [27]. In medical imaging, GANs play a crucial role in super-resolution, where they enhance image clarity and detail, and in the generation of synthetic images. The creation of these synthetic images is fundamental for data augmentation, training simulations, and the provision of enhanced diagnostic insights without the need for additional radiation exposure or patient involvement [28] [15]. Traditionally, GANs have predominantly utilized CNNs for both the generator and discriminator. However, the emergence of ViTs has led researchers to investigate more efficient architectures for ViT-based GAN models [29] [30]. ViTs are particularly effective in brain imaging, excelling at capturing comprehensive brain patterns, thus ensuring a more complete representation and superior feature extraction. This capability is especially beneficial in recognizing complex neural structures, surpassing the performance of CNNs. For instance, [31] introduced a pre-trained ViT model for classifying brain tumors, addressing the limitations of CNNs that tend to focus predominantly on minute pixel variations. Additionally, [32] demonstrated an enhanced ViT architecture capable of utilizing both structural and functional MRI data for predicting various stages of Alzheimer’s disease. Furthermore, the integration of ViT and GAN has emerged as a novel trend in medical imaging. An example of this is the study by Zhao et al. [33], who developed a Swin Transformer-based GAN model [34] aimed at effective reconstruction of high-resolution MRI images.

In the domain of medical image synthesis, the focus has been on generating images across different modalities, such as CT, MRI, PET, and others. Dalmaz et al. [35] created a new GAN model that combines CNNs with transformer blocks. This model makes it much easier to make medical images that are similar and work better. However, we were hardly able to find related works that corresponded to MRI structural and functional image synthesis, besides our previous works, which synthesized FNC data from given sMRI and achieved a high correlation between real FNC and generated FNC data [36]. However, the previous work was using a basic ViT-based GAN architecture, which is time-consuming. In addition, we didn’t generate structural biomarkers, which is also a shortcoming.

## III. Methods

Our methodology is a cutting-edge combination of deep learning architectures geared to the complex problem of generating functional neural connectivity (FNC) maps from massive amounts of three-dimensional sMRI data. A ViT-based conditional generative adversarial network (cViT-GAN) is at the heart of our technology, and it is meant to learn and create high-fidelity FNC representations. We propose an efficient block-wise self-attention technique to avoid the significant computational overhead generally experienced by ViT’s processing of tiny picture patches. This personalized strategy retains the ViT’s tremendous feature extraction capabilities while preserving computational efficiency, allowing the model to manage the large amounts of data associated with sMRI. We extend our methods by utilizing the attention weights produced from each layer of the ViT encoders. When these weights are superimposed onto the spatial information recorded by 3D MRI images, they make it easier to create biomarkers. These are not only activations but sophisticated attention maps reflecting the different brain patterns that distinguish schizophrenia (SZ) and healthy control (HC) participants. Our technique paves the door for more insightful neuroimaging investigations, possibly assisting in early diagnosis and intervention efforts for mental health issues, by giving a visual and quantitative differentiation between the two groups.

### A. Generative Adversarial Networks

Integrating generative adversarial networks (GANs) into the domain of medical imaging necessitates a nuanced understanding of their loss functions. For our specific application of synthesizing FNC maps from three-dimensional sMRI data, we construct a composite loss function that ensures the generation of realistic and medically informative images. The total loss ℒ _*total*_ of our GAN framework is a weighted sum of four components:

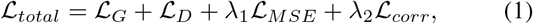

where ℒ _*G*_ denotes the generator loss, ℒ _*D*_ the discriminator loss, ℒ_*MSE*_ the mean squared error loss, and ℒ_*corr*_ the correlation loss. The terms *λ*_1_ and *λ*_2_ are hyperparameters that balance the contribution of the MSE loss and the correlation loss, respectively.

The generator loss ℒ_*G*_ is defined as:

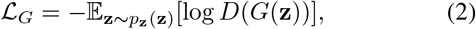

where *G* is the generator, *D* is the discriminator, and **z** is a point sampled from the generator’s input noise distribution *p*_**z**_(**z**). The discriminator loss *ℒ* _*D*_ is formulated as:

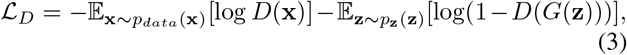

where **x** represents real data samples from the distribution *p*_*data*_(**x**). The mean squared error loss ℒ _*MSE*_ is incorporated to penalize the pixel-wise differences between the generated and real images, thus preserving the structural integrity of the FNC maps:

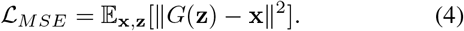

The innovation in our approach is embodied by the correlation loss ℒ _*corr*_, which ensures that the statistical dependencies between regions in the generated FNC maps are reflective of the true data. This is crucial for maintaining the biological fidelity of the neural connectivity patterns. The correlation loss is defined as:

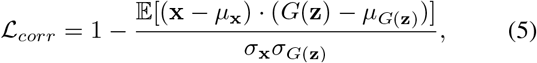

where *μ* and *σ* denote the mean and standard deviation, respectively. This loss encourages the generated maps to have a correlation structure similar to that of the real FNC maps.

Our GAN architecture also incorporates a conditional input, whereby the generator receives both a sample of noise **z** and a label indicating the class (SZ or HC). This guides the generator towards producing FNC maps that are not only realistic but also correctly aligned with the specified condition:

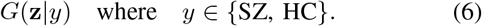

In essence, by carefully crafting the loss function and incorporating conditionality, our method aims to drive the GAN towards producing medically valuable outputs, tailored to the requirements of the task at hand.

### B. Vision Transformer

The vision transformer (ViT) is an innovative neural network architecture that adapts the mechanisms of transformers, originally designed for natural language processing, to the domain of computer vision. ViT’s core idea is to treat image pixels as a sequence of tokens, akin to words in a sentence, and apply self-attention mechanisms to capture global dependencies within the image.

#### 1) General Architecture

Incorporating ViT within our GAN architecture leverages its powerful representational capabilities for both the generator and the discriminator, enabling the processing of complex three-dimensional MRI data. The generator employs a 3D ViT architecture that begins by dissecting the 3D MRI input into a series of 3D small patches. These patches are analogous to tokens in a transformer model. Each patch is then flattened and passed through a series of linear embedding layers to convert it into a suitable format for the transformer encoder:

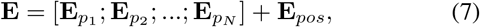

where 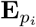 represents the embedded representation of the *i*-th patch, *N* is the total number of patches, and **E**_*pos*_ denotes the positional encodings added to retain the notion of the order of patches. These embeddings serve as the input to the ViT encoder, which comprises multiple layers of multi-headed self-attention and feed-forward networks:

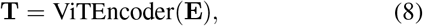

where **T** denotes the sequence of transformer encoder outputs corresponding to each patch embedding. Subsequently, each token produced by the ViT encoder is passed through a multilayer perceptron (MLP) network. This MLP is designed to reconstruct the small patches of the generated FNC matrix, transforming the abstract representations learned by the ViT into spatially structured outputs:

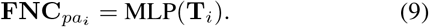

The collection of FNC patches 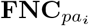 is then reassem-bled to form the complete FNC map, which serves as the generator’s final output:

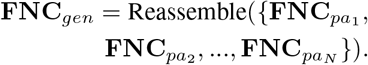

For the discriminator, the 3D ViT discerns between the real and generated FNC maps, employing a similar patch-based approach to extract features and perform classification. The discriminator’s role is to evaluate the authenticity and quality of the generated FNC maps, guiding the generator through the adversarial training process to produce outputs that are increasingly indistinguishable from the real FNC maps derived from sMRI data. By integrating the ViT model into both the generator and discriminator of our GAN, we harness its potent capacity for capturing intricate patterns and dependencies within the complex data structure of three-dimensional brain imaging.

#### 2) Pre-trained 3D Patch Embedding

The utility of pretrained models in deep learning is unparalleled, particularly in domains where data is scarce or where training from scratch is computationally prohibitive. Leveraging a pre-trained 3D ViT model, our generator benefits from an advanced starting point. This model, initially trained on upstream tasks such as the classification of SZ and HC from MRI data, has already learned a rich hierarchy of features that are highly relevant to our target domain. The pre-trained model forms the cornerstone of our generator’s architecture. Specifically, for the patch embedding process, we utilize the pre-trained embeddings, denoted as:

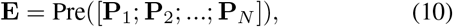

where **P**_*i*_ corresponds to the flattened vector of the *i*-th 3D patch, and *N* is the number of non-overlapping 3D patches extracted from the sMRI input. The function PretrainedEmbed (·) encapsulates the process of obtaining the embedded representations using the pre-trained ViT model.

These pre-trained patch embeddings already encode the spatial hierarchies learned from the upstream classification task, providing a richly structured feature space that is fine-tuned for the generator:

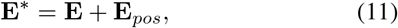

where **E**^*∗*^ represents the embeddings that will be utilized in the transformer encoder, and **E**_*pos*_ is the positional encoding added to the pre-trained embeddings.

The integration of these pre-trained embeddings allows our generator to rapidly adapt to the generation of FNC maps, leveraging the already learned discriminative features between SZ and HC groups. This not only accelerates the training process but also imbues the downstream task with a depth of learned representation that would be otherwise unattainable in a reasonable time frame:

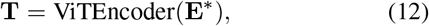

where **T** signifies the sequence of transformer encoder outputs based on the pre-trained embeddings, propelling our GAN towards efficiently generating meaningful and accurate FNC maps. This pre-trained 3D ViT embedding method enhances the generator’s capability, facilitating a more effective transfer of knowledge from the upstream task to the downstream synthesis of FNC matrices, establishing a methodological synergy that underpins the sophistication of our model.

#### 3) Block-wised Multi-head Self-attention

Incorporating the block-wise multi-head self-attention (BMHSA) [37] mechanism into our model optimizes computing efficiency while keeping the delicate features required for high-resolution biomarker detection from 3D MRI data. We used BMHSA in vision tasks because of its excellent performance in dealing with long-text in NLP tasks. BMHSA separates the collection of 3D MRI patch embeddings into smaller, more manageable chunks, enabling for more focused and computationally efficient self-attention within these partitions. This is especially useful for dealing with the small patch sizes characteristic of 3D MRI data, which are necessary to preserve the resolution required for proper biomarker analysis. Within each block, BMHSA operates by computing self-attention independently, which drastically reduces the overall computational load compared to traditional methods. Mathematically, the self-attention within a block *b* can be expressed as:

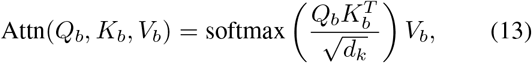

where *Q*_*b*_, *K*_*b*_, and *V*_*b*_ are the queries, keys, and values for the block *b*, and *d*_*k*_ represents the scaling factor for the dot products within the softmax function to ensure numerical stability.

Leveraging the concept of multi-head attention, BMHSA allows the model to concurrently attend to different representational subspaces and positions within each block, formulated as:

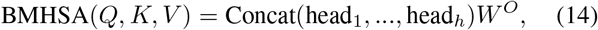

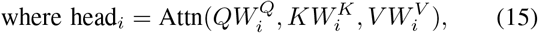

with each 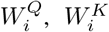, and 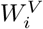 denoting the respective parameter matrices for each attention head *i*, and *W*^*O*^ being the output linear transformation matrix.

The BMHSA approach ensures the emphasis of intra-block (regional) relationships while facilitating the preservation of inter-block (long-range) dependencies. These long-range dependencies are crucial for the analysis of structural brain images, as they allow the model to piece together localized information to form a comprehensive understanding of the brain’s structure:

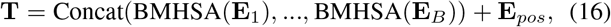

In this equation, **T** is the output of all the transformer encoder layers put together. It includes both detailed and general information about the brain’s structure. The **E**_*B*_ terms show the embeddings from each block, and the **E**_*pos*_ terms show the positional encodings that are needed to keep the 3D MRI data’s natural spatial relationships.

##### BMHSA Complexity Analysis

By employing the block-wise multi-head self-attention (BMHSA) mechanism, our model achieves significant reductions in computational costs while successfully generating high-resolution attention maps. Traditional self-attention mechanisms, such as those used in ViT models, exhibit a computational complexity that scales quadratically with the length of the sequence *n*. This complexity is expressed as *O*(*n*^2^ · *d*), where *d* is the dimensionality of the attention heads. For long sequences (such as tokens or patches), this scaling becomes computationally prohibitive.

BMHSA addresses this issue by partitioning the input sequence into smaller, fixed-size b locks, e ach o f l ength *k*. Within each block, self-attention is computed independently, leading to a complexity of *O*(*k*^2^ · *d*) per block. If the input sequence is divided into *m* such blocks, with the total sequence length *n* being equal to *m* × *k*, the initial thought would be to express the overall complexity as the sum across all blocks, leading to *O*(*m* · *k*^2^ · *d*).

However, a more accurate representation of BMHSA’s complexity takes into account the parallelizability of these block computations. Since each block’s computation is independent, the per-block complexity of *O*(*k*^2^ · *d*) remains, but the computations across different blocks can be performed in parallel. Therefore, the overall computational load does not directly scale with the number of blocks *m*.

Thus, the total computational complexity of BMHSA can be more accurately described as:

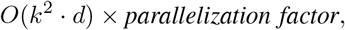

In conclusion, by judiciously choosing an appropriate block size *k*, BMHSA effectively balances the trade-off between manageable computational costs and the granularity of attention required for detailed analysis in tasks such as highresolution biomarker detection from 3D MRI data.

### C. cEViT-GAN Architecture

The cEViT-GAN architecture is a novel paradigm designed exclusively for the study of 3D sMRI data, with a focus on the synthesis of sFNC maps. This paradigm is very important in the context of neuroscientific research, particularly in the investigation and understanding of neurological illnesses such as schizophrenia. The cEViT-GAN, in contrast to standard convolutional techniques, employs a purely self-attention mechanism. This distinguishes it as a purely ViT-based GAN model, as opposed to traditional CNN-based GAN architectures. Such a design choice emphasizes the device’s originality in the field of medical picture processing and analysis. Table 1 details the many layers and functions in our cEViT-GAN model. Figure 2 depicts the pipeline and overall architecture of cEViT-GAN.

**TABLE 1.** cEViT-GAN Architecture Overview.

| Component | Layer/Function | Description |
| --- | --- | --- |
| Generator | Input | Processes 3D sMRI patches labeled SZ or HC |
|  | Pre-trained 3D Embedding | Utilizes pre-trained ViT model for feature extraction |
|  | BMHSA Layers | Manages computational load, extracts features |
|  | MLPs for Reconstruction | Converts encoder outputs to structured FNC patches |
|  | Output Assembly | Assembles patches into complete FNC map |
| Discriminator | Patch Embedding | Divides FNC maps into patches for processing |
|  | ViT Encoder | Processes embedded patches, extracts features |
|  | Classification Output | MLP that outputs probability of real or generated map |

**Fig. 1.**
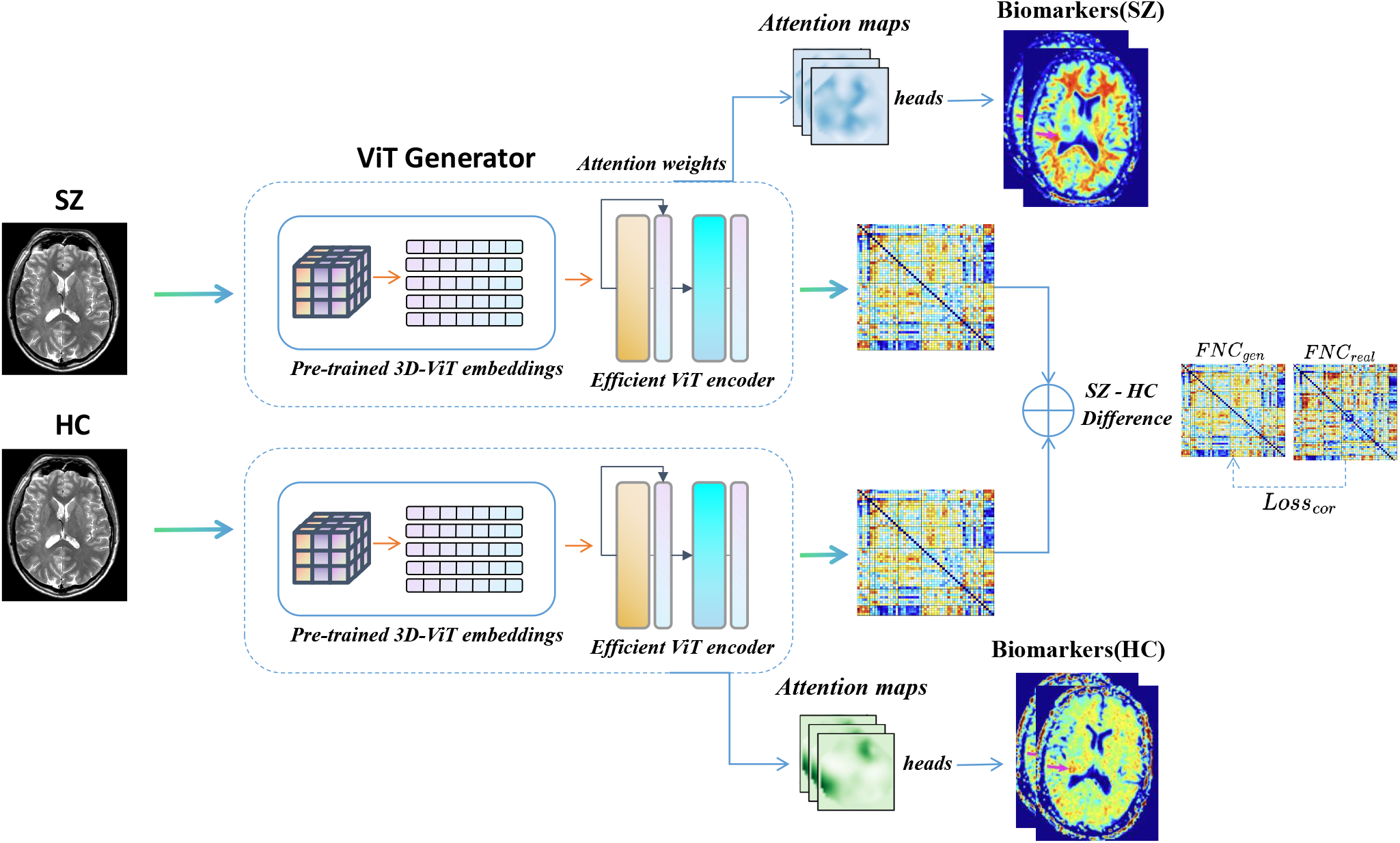
The proposed methodology involves the analysis of brain MRI scans, specifically those labeled as SZ and HC. The objective is to generate group difference FNC data by utilizing an efficient generator from the cViT-GAN framework. Additionally, attention weights are extracted from the ViT encoder to obtain 3D MRI attention maps for the different groups. We then apply this approach to identify biomarkers associated with schizophrenia.

**Fig. 2.**
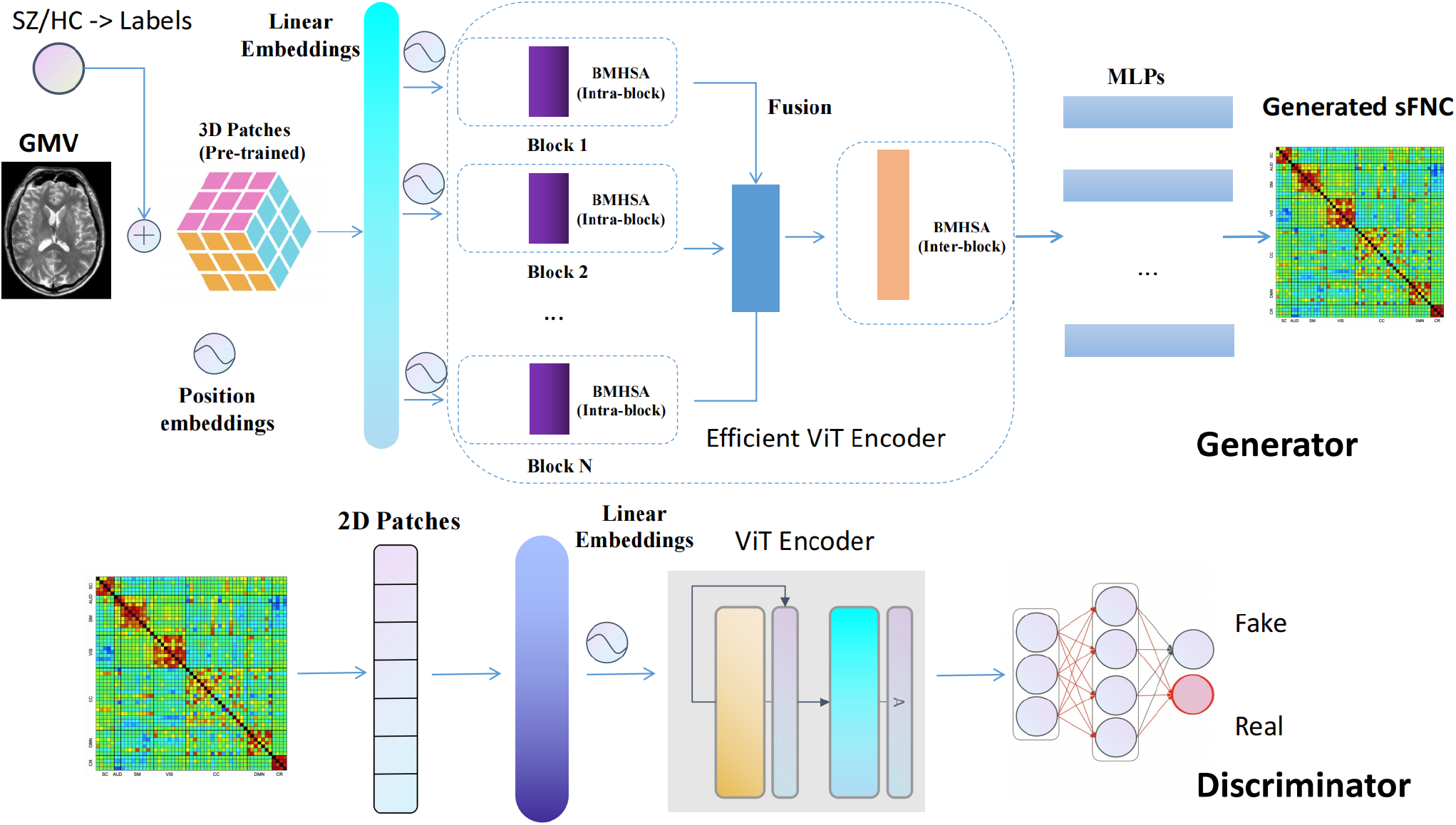
cEViT-GAN’s detailed architecture: The 3D GMV and its label (SZ/HC) were fed into the generator, which then passed through the pre-trained 3D embedding layers before entering the efficient ViT encoder layer. Following that, a series of MLP outputs are used to form the resulting sFNC. The architecture of the discriminator is fairly similar to that of a typical 2D ViT.

#### Generator Architecture

The generator begins by taking small 3D sMRI patches, labeled as either SZ or HC. These patches are initially processed through pre-trained 3D embedding layers, utilizing the pre-trained ViT model to capitalize on its extensive feature extraction capabilities from medical imaging data. The data then passes through BMHSABMHSA layers, which are crucial for efficient feature extraction and computational load management. The final stage involves MLPs reconstructing the sFNC maps from these features, converting transformer outputs into spatially structured FNC patches, which are then assembled into a complete FNC map.

#### Discriminator Architecture

The discriminator’s design parallels a 3D ViT. It begins by dividing 3D FNC maps into patches, which are then processed through the ViT encoder. This enables effective discernment of patterns within the FNC maps. The discriminator ultimately classifies the input based on these features, producing a probability score indicating whether the FNC map is real or generated. This feedback is vital for the adversarial training of the generator, aiming for the creation of accurate and realistic FNC maps.

## IV. Experiment

This part will talk about how we set up our experiment (datasets and preprocessing), how we trained and tested the models (including setting baselines), the special cEViT-GANs and their settings, and how we did our experiments to see how the brain works structurally and functionally.

### A. Experimental Setups

#### 1) Datasets

In our study, we utilized two comprehensive datasets pertinent to clinical schizophrenia research. Dataset 1 amalgamated data from three distinct studies: fBIRN (Functional Imaging Biomedical Informatics Research Network) across seven sites, MPRC (Maryland Psychiatric Research Center) spanning three sites, and COBRE (Center for Biomedical Research Excellence) at a single site. This aggregation culminated in a total of 827 participants, comprising 477 control subjects (average age: 38.76 ± 13.39, encompassing 213 females and 264 males) and 350 individuals diagnosed with schizophrenia (average age: 38.70 ± 13.14, including 96 females and 254 males). The fBIRN dataset was acquired using uniform resting-state fMRI (rsfMRI) parameters across all sites. We used a standard gradient echo-planar imaging (EPI) sequence with a repetition time (TR) of 2000 ms and an echo time (TE) of 30 ms. The voxels were 3.4375 × 3.4375 × 4 mm in size, and the field of view (FOV) was 220 × 220 mm. The data was captured using six Siemens Tim Trio 3-Tesla scanners and one General Electric Discovery MR750 3.0 Tesla scanner. In the COBRE segment, rsfMRI images were also taken using a standard EPI sequence, but with a slightly different TR/TE of 2000/29 ms and voxel sizes of 3.75 × 3.75 × 4.5 mm, within a field of view (FOV) of 240 × 240 mm, using a 3-Tesla Siemens Tim Trio scanner. The MPRC dataset was gathered using a trio of distinct 3-Tesla Siemens scanners, namely the Siemens Allegra, Trio, and Tim Trio.

Dataset 2 encompassed data from 815 participants hailing from seven distinguished Chinese hospitals: Peking University Sixth Hospital, Beijing Huilongguan Hospital, Xinxiang Hospital Simens, Xinxiang Hospital GE, Xijing Hospital, Renmin Hospital of Wuhan University, and Zhumadian Psychiatric Hospital. This dataset included 326 control subjects (average age: 29.81 ± 8.68, with a gender distribution of 167 females and 159 males) and 489 individuals with schizophrenia (average age: 28.98 ± 7.63, consisting of 229 females and 260 males), all of Han Chinese descent. The resting-state fMRI data from these participants were collected using three different 3-Tesla scanners across the sites, namely the Siemens Tim Trio, Siemens Verio, and Signa HDx GE Scanner. Participants were instructed to maintain stillness, relaxation, and wakefulness during the scans.

#### 2) Pre-processing

To prepare the fMRI data, several critical processes were required: slice timing correction, realignment, normalization to the EPI template, and smoothing with a 6 mm kernel. Our prior studies contain detailed descriptions of these preprocessing methods. Furthermore, sFNC data was obtained using fMRI time series cross-correlation analysis. As spatial priors, a fully automated spatially limited ICA method and the NeuroMark template [38] were utilized. We used a voxel-based morphometry process on the sMRI data to acquire voxel-level gray matter volume data.

### B. Models

### C. Baselines

The primary goal of our comprehensive investigation of the efficacy and performance of several GAN models was to evaluate these models in terms of image-generating capabilities and output quality. The baseline models for comparison were carefully chosen, with special consideration given to their relevance to our pioneering work in synthesizing sFNC from sMRI. While there are no clear antecedents for synthesizing FNC, the closest similarity is found in the realm of picture synthesis. As a result, we chose GAN models known for their expertise in this field as our baselines.

The first group of baselines includes CNN-based GAN models like Pix2Pix and deep convolutional GAN (DCGAN). The Pix2Pix model, which employs a U-Net generator and a PatchGAN discriminator, is well-known for its ability to solve image-to-image translation problems. The importance of this model in our research arises from its demonstrated ability to generate high-fidelity images from input photographs, a process that is similar to our goal of sFNC synthesis from sMRI data. The integration of low-level and high-level characteristics in the generator by the U-Net architecture improves the detail and quality of the output images. Furthermore, the PatchGAN discriminator focuses on judging the realism of local image patches, which contributes greatly to image sharpness and overall coherence. As a result, these models provide a solid foundation for assessing the potential of GANs in our ground-breaking effort to synthesize sFNC from sMRI. Moreover, we use traditional self-attention-based cViT-GAN as other baselines, which can show the efficiency of our model.

### D. cEViT-GANs

Our research into novel GAN models resulted in the creation of the cEViT-GAN framework, a revolutionary technique developed exclusively for sFNC synthesis from sMRI. The cEViT-GAN models incorporate cutting-edge approaches, including a pre-training strategy focused on embedding 3D patches and efficient usage of ViT blocks via blockwise self-attention. This novel combination intends to improve picture synthesis quality by combining the strengths of CNNs and transformers.

To fine-tune cEViT-GAN’s performance for our unique application, we created two primary modifications. The first form, cEViT-GAN-b3, is made up of three parallel blockwise multi-head self-attention (BMHSA) blocks. This setup does not include interblock self-attention, allowing the model to be optimized for speed and efficiency while still producing high-quality images. The cEViT-GAN-b3large variant expands on the cEViT-GAN-b3 design by including an interblock self-attention mechanism. This update aims to increase the model’s ability to capture and integrate more complicated patterns and relationships in data, potentially leading to more accurate and detailed sFNC synthesis.

Furthermore, we have introduced cEViT-GAN-b6, which makes use of six parallel blocks in the BMHSA layer. It, like cEViT-GAN-b3, avoids interblocking self-attention. This variant is intended to evaluate the impact of increasing the number of parallel blocks without the complexity of interblock connections, providing an alternate method to balancing computational efficiency and picture synthesis quality.

Our core model, cEViT-GAN-b3large, serves as the foundation of our investigation. This model serves as the foundation for all visualizations and analyses in our study. The cEViT-GAN-b3large architecture is our most advanced attempt at using the promise of GANs for sFNC synthesis. It has a mix of several BMHSA layers that interblock self-attention. Its design attempts to achieve an optimal balance between capturing precise, high-level features and ensuring efficient 3D sMRI data processing. The addition of interblock self-attention is especially noteworthy since it allows the model to more effectively integrate information across multiple layers, potentially leading to a more nuanced and accurate synthesis of sFNC from sMRI data.

### E. Experiments Details

#### 1) Pre-training

When we test all of our ViT-based GAN models, including the baseline models and our cViT-GAN variants, we use 3D linear embeddings that have already been trained. These embeddings come from a multimodal deep learning model we made in earlier work that was mostly about putting people with schizophrenia into groups [17]. Our earlier model and the 3D ViT embedding layer used in this study are similar in how they are built, which gives our GAN models a strong base.

This pre-training step is particularly significant for the generator components of our ViT-based GANs. Our cEViT-GANs’ generators have a deep understanding of complex patterns in medical imaging data from the very beginning because they start with the weights from these pre-trained 3D linear embeddings. This approach not only accelerates the training process by providing a well-informed starting point but also enhances the overall efficiency and effectiveness of the models. The pre-trained embeddings encapsulate a rich array of features relevant to SZ, a complexity that is beneficial for our current task of sFNC synthesis from sMRI data. The embeddings carry nuances of neuroimaging patterns associated with SZ, offering a unique perspective that is particularly relevant given the neurological focus of our study. When applied to our GAN models’ generator networks, these embeddings make it possible for the generators to make sFNC images that are more nuanced, detailed, and clinically relevant.

#### 2) Train and Validation

It’s worth noting that all CNN-based GAN models use the same set of parameters and training techniques, whereas ViT-based GANs (including baseline and cEViT-GAN variations) use a different set.

We use Kaiming initialization to select initial weights for CNN-based GANs. We use the AdamW optimizer for both the generator and the discriminator and set the learning rate to 1e-3. The learning schedule employs MultistepLR, with adjustments occurring at the 20th, 50th, and 150th epochs. Due to the unique nature of ViT training, ViT-based GAN models require a lower learning rate. In all baseline and cViT-GAN variations, we use pre-trained weights for the 3D patch embedding components of the GAN’s generator. The learning rate is 1e-4 in this case, with AdamW serving as the optimizer for both the generator and the discriminator. The training plan is MultistepLR once more, but this time with changes at the 20th, 50th, and 90th epochs. The pre-training stage facilitates convergence around the 90th epoch in this timetable.

Using cross-validation on both types of models improves model resilience and dependability. This technique is useful for determining how the models would perform on different data sets, decreasing the danger of over-fitting and assuring generalization. Our training and validation operations are powered by 8 NVIDIA Tesla V100 GPUs. Furthermore, we use parallel and distributed training approaches to spread the training strain across numerous GPUs. This method dramatically improves processing efficiency and shortens training times. It’s especially useful for dealing with the massive amounts of data and sophisticated neural network structures involved in our research.

### F. Evaluation Metrics

In our research, we deploy a trio of critical metrics to assess the efficacy of our model in synthesizing Functional Network Connectivity (FNC) patterns. These metrics include the Mean Squared Error (MSE), the Pearson Correlation Coefficient, and the Cosine Similarity. Each of these metrics plays a crucial role in evaluating the precision and reliability of the FNC patterns generated by our model, offering distinct insights into the model’s performance and facilitating a comprehensive assessment when compared to authentic FNC data.

#### 1) Mean Squared Error (MSE)

MSE is a metric commonly utilized in regression analysis and signal processing. It quantifies the average of the squares of errors, which are the differences between the estimated values and the actual values. In the context of our FNC data, the MSE is calculated as follows:

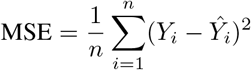

Here, *n* denotes the total number of FNC entries, *Y*_*i*_ represents the actual FNC value, and *Ŷ*_*i*_ signifies the estimated FNC value produced by the model. A lower MSE value is indicative of superior model performance, signifying a reduced deviation from the true FNC values.

#### 2) Pearson Correlation Coefficient

The Pearson Correlation Coefficient is a measure that quantifies the linear correlation between two datasets. It yields a value within the range of −1 to 1, where 1 denotes a total positive linear correlation, 0 signifies no linear correlation, and −1 indicates a total negative linear correlation. In evaluating our model, this coefficient is defined as:

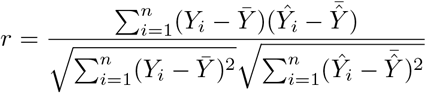

Where 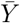 and 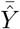 are the mean values of the actual and estimated FNCs, respectively. A higher absolute value of this coefficient implies a stronger correlation between the generated FNCs and the real data.

#### 3) Cosine Similarity

Cosine Similarity is a metric employed to ascertain the similarity between two vectors, irrespective of their magnitude, and is especially pertinent in high-dimensional spaces. The cosine similarity between the actual and the model-generated FNC vectors is computed as:

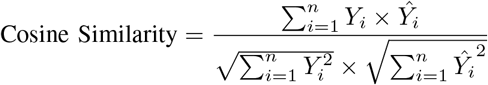

In this formula, the numerator represents the dot product of the actual and estimated FNC vectors, while the denominator is the product of the Euclidean norms of these vectors.

Together, these metrics provide a strong and flexible way to check how accurate and similar the FNC patterns our model creates are to real FNC data. This gives us important information about how well the model can copy complex neural connectivity patterns.

### G. Visualizations

We performed intense brain anatomical and functional visualization based on self-attention operations in two phases. The first step is to use attention weights on original brain maps to find any biomarkers in MRI data while also making a matching FNC. The second step is to compare the made FNC with the real FNC, which could help find functional biomarkers for SZ disease.

#### 1) MRI Attention Maps

To extract attention weights, we used a rollout method. We concatenated the weights from each block to achieve blockwise multihead self-attention. We superimposed the weights of the interblock self-attention onto the averaged block weights when using interblock self-attention. This method produced a thorough attention map. This approach is useful for detecting possible biomarkers in brain MRI data. High attention weights help us identify regions of interest that may be connected with various neurological diseases, such as SZ. The attention map serves as a guide, showing the most important sections of the brain MRI data. Researchers can get insights into the fundamental mechanisms of SZ and potentially other neurological illnesses by better understanding these areas. The incorporation of attention weights into MRI data analysis represents a significant improvement in neuroimaging and the research of brain diseases. Fortunately, analyzing the group difference (HZ-HC) attention map allowed us to identify brain areas strongly associated with SZ that aligned with our existing knowledge.

#### 2) FNC Maps

Our cEViT-GANs generate reasonably accurate FNC maps. Researchers can use these maps to supplement current data and identify biomarkers that fill gaps in structural and functional data. We generate FNCs for each sMRI and then average the FNC maps based on the labels (SZ or HC). We also evaluate group differences in FNC maps to depict the distinctive link between the two groups.

## V. Results

This section presents the outcomes of the experiments as well as a visualization of the structure and function of the brain using attention maps and FNC biomarkers. Initially, we conducted exhaustive experiments on various baselines and our cEViT-GAN variants to ensure that our model exhibited superior accuracy and robustness.

### A. Model Performance

We did a series of experiments that can compare the model performance of our baseline models (Pix2Pix, DCGAN, and cViT-GAN) and our cEViT-GAN and its variants. Figure 3 shows the basic model performance between these baselines and our new models.

**Fig. 3.**
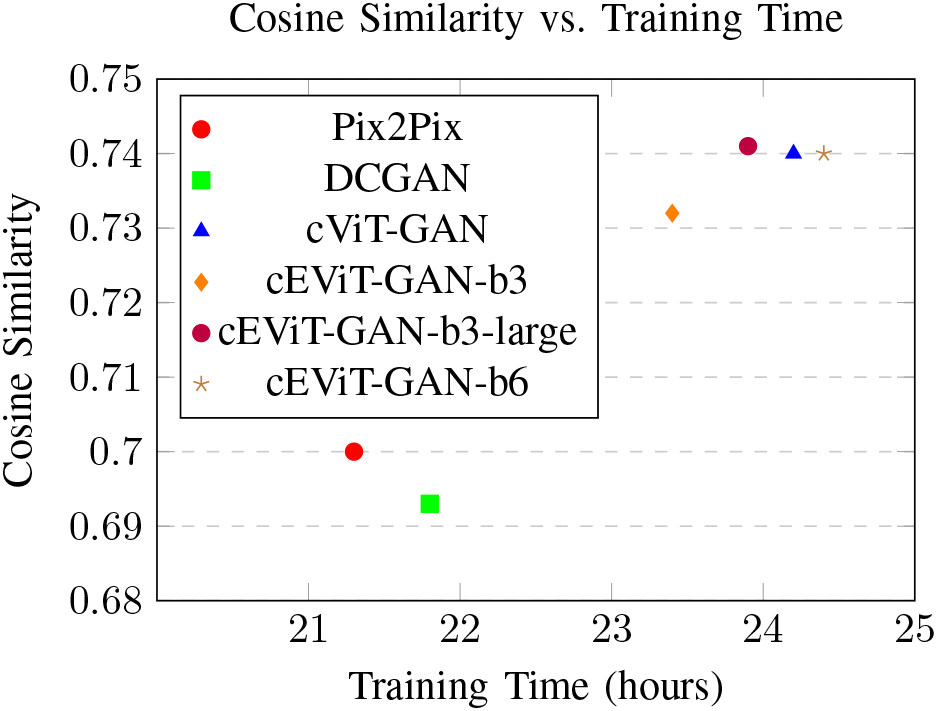
Comparison of different models in terms of Cosine Similarity and Training Time.

### B. MRI Attention Maps

We analyzed attention weights in our cEViT-GAN generator to generate our 3D MRI attention maps. We created subject-specific attention maps for each member of our testing set, then tested for group differences using a two-sample t-test. Each voxel in our attention maps represents a t-value from this statistical test. To account for multiple comparisons, we used the false discovery rate (FDR) method with a *q <* 0.05 threshold. This approach accounts for the possibility of type I errors while running several statistical tests. The attention maps that arise emphasize areas with statistically significant changes in activation patterns between groups. Figure 4 shows the attention maps in a three-plane view, including a rendering.

**Fig. 4.**
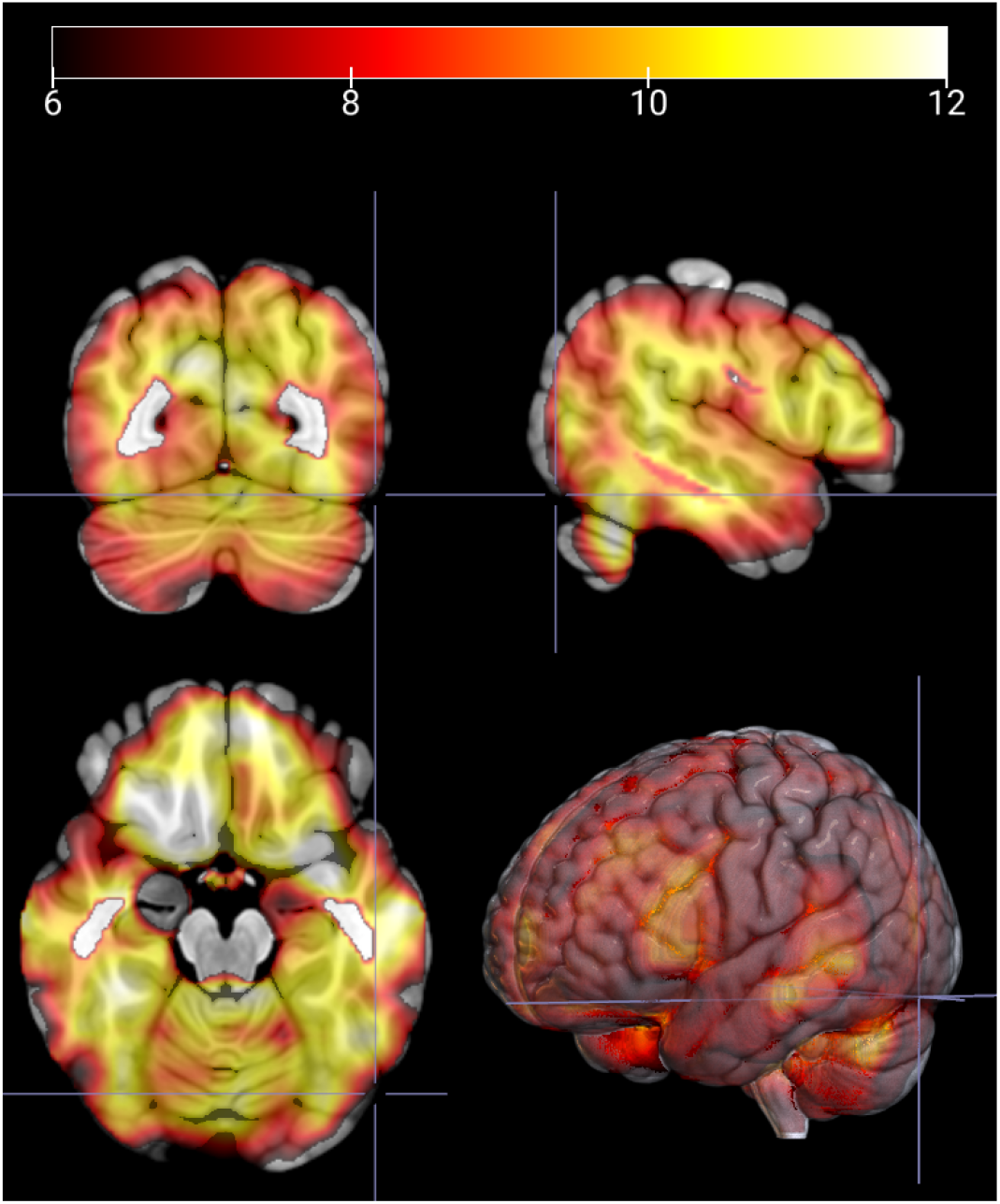
The 3D MRI attention maps for group difference analysis (SZ vs HC), which indicate the significant ROIs that are strongly associated with schizophrenia disease.

The results clearly highlight structural regions know to be im-plicated in schizophrenia including temporal lobe, cerebellum, and frontal lobe.

Figure 4 indicates that while generating the related functional outputs, our model prioritized four brain regions: the medial pre-frontal cortex (mPFC), the dorsolateral prefrontal cortex (DL-PFC), the temporal lobe, and the cerebellum. Schizophrenia is a diverse, complex psychiatric condition that frequently involves dysfunctions in numerous brain circuits. Based on traditional neuroscience and previous knowledge, mFPC is intimately related to executive processes and decision-making, both of which can be affected in schizophrenia [39]. The mPFC is also involved in emotional processing, and abnormalities here can be linked to negative schizophrenia symptoms including apathy and social disengagement [40]. DL-PFC is required for cognitive control and working memory, both of which are frequently impaired in people with schizophrenia. Deficits in this area can contribute to the disorder’s hallmarks of disorganized thinking and trouble focusing attention. The superior temporal gyrus, in particular, is connected with auditory processing and language. Temporal lobe dysfunction has been linked to auditory hallucinations and language difficulties seen in schizophrenia patients [41]. Finally, the cerebellum’s significance in cognitive processing is now recognized. Cerebellar abnormalities may contribute to cognitive impairments and affective dysregulation in schizophrenia, according to recent research [42], [43].

### C. FNC Biomarkers

#### 1) FNC Analysis

In this study, a sophisticated GAN model was employed to generate FNC outputs from a test dataset. Our analysis revealed that the model’s output for the whole average FNC exhibited a strong correlation (0.97) with the actual FNC data across all subjects. This is effectively visualized in Figure 5, which compares the model-generated whole average FNC with the genuine FNC data.

**Fig. 5.**
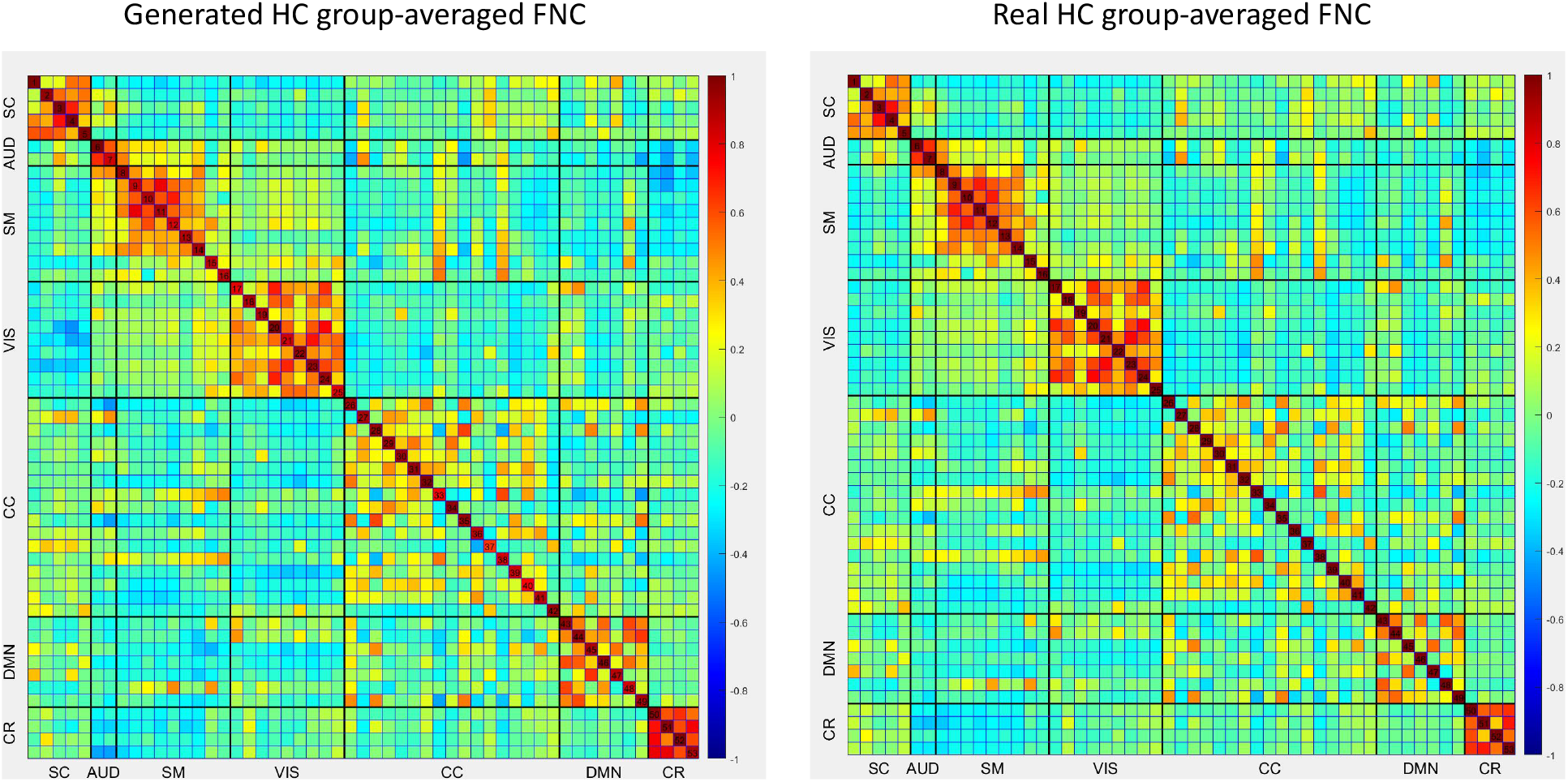
The generated whole average FNC vs. real whole average FNC

**Fig. 6.**
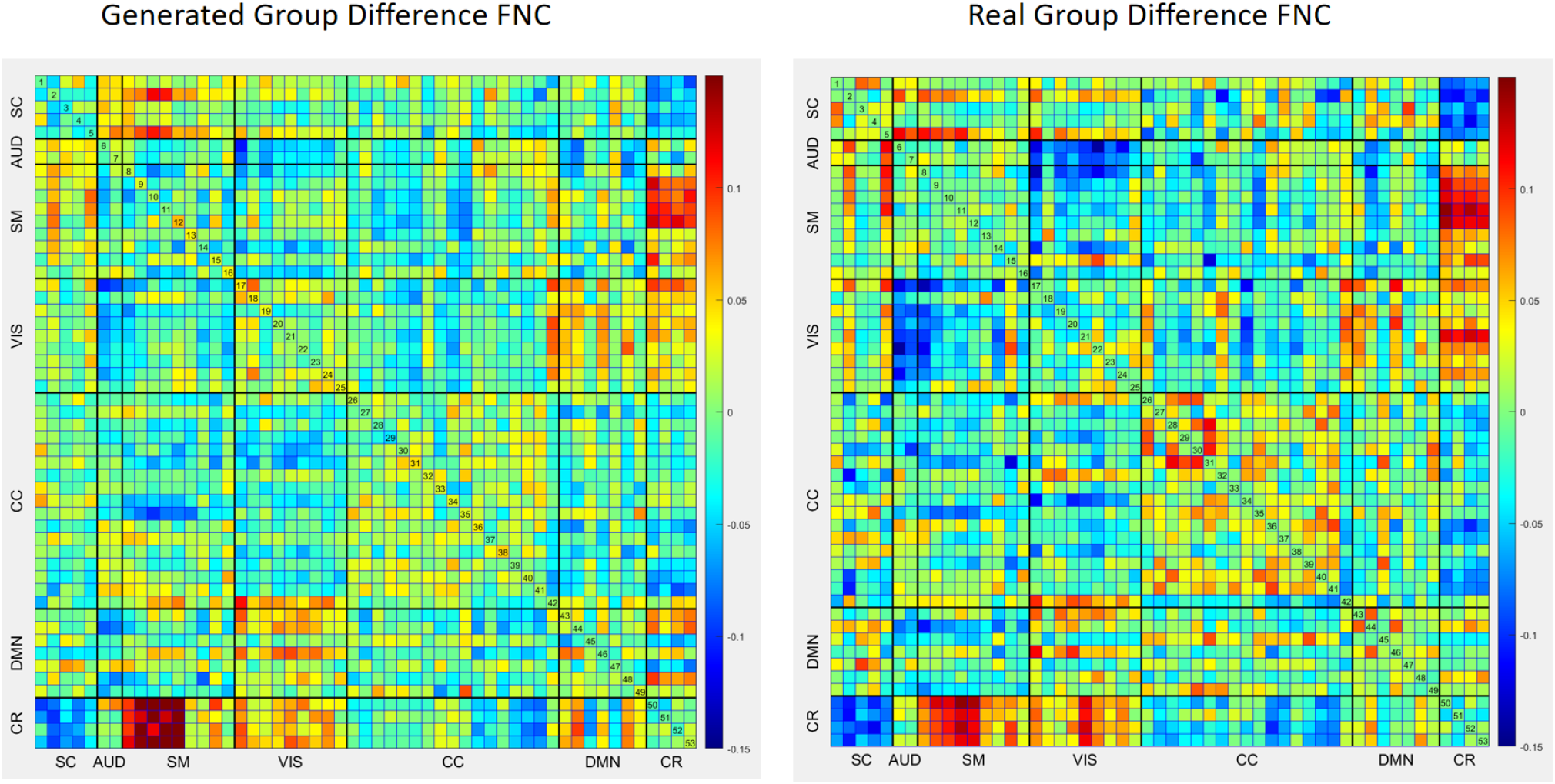
The generated ‘fake’ group difference FNC matrix vs. real group difference FNC matrix.

The ability of our GAN model to replicate FNC from 3D MRI scans of gray matter with high accuracy can be credited to the identification of neural structures via independent component analysis (ICA). ICA has been known to uncover network-like structures within the resting gray matter, providing insights into the model’s capability to replicate these intricate neural patterns. The correlation observed in our model’s output with the real data not only validates our approach but also aligns it with previous scientific research in neuroimaging [44], [16], [11].

#### 2) Group Difference Analysis

We also show the produced and genuine group-difference FNC (HC-SZ). Figure 5 shows a comparison of calculated and actual FNC group differences. Our model can infer group-difference FNC from brain structure with a remarkably high correlation (0.74) especially given brain function contains unique information above and beyond brain structure. There are also some ways we can further explore approaches to further improve the prediction. Individual FNC patterns within each group (SZ or HC) are likely more constant, making high-fidelity replication easier for the GAN. However, when comparing groups, the model may lose nuanced, individual-specific functional information that is critical in discriminating between SZ and HC on a more granular level.

However, our cEViT-GAN model can identify a strong similarity between the generated group-difference FNC and the real one, and the patterns are those that are know to be implicated in schizophrenia, including subcortical areas. These include connections between the cerebellum and the subcortical (CB-SC), auditory (CB-AUD), somatomotor (CB-SM), visual (CB-VS), cingulo-opercular (CB-CC), default mode (CB-DM), and the cerebellum itself (CB-CB). The synthetic FNC data obtained by structural MRI has a remarkable correlation with real FNC data, with similarities reaching 0.85 in certain subcortical linkages.

This important finding shows that subcortical structures are important for identifying differences between HC and SZ participants and that the cEViT-GAN model does a good job of showing these important structural-functional connections. The remarkable similarity in subcortical areas demonstrates that our model is quite good at replicating complicated, potentially clinically relevant, brain patterns. Such skills signal new opportunities for improving our understanding of illnesses such as schizophrenia, offering more precise diagnostic measures, and personalizing therapy methods. Furthermore, our model demonstrates that there is a high level of agreement in the difference in values for other pairs of connections, such as cingulo-opercular (CC-CC), somatomotor-default mode (SM-DM), and visual-default mode (VS-DM). We also find moderate parallelism in visual-auditory (VS-AUD) and cingulo-opercular-somatomotor (CC-SM) pairs. These findings provide greater insight into how the disparities in FNC observed between the HC and SZ groups may be caused by underlying structural issues. The combined insights are critical for developing more refined diagnostic tools and therapy approaches for navigating the complexities of schizophrenia.

#### 3) Cross-domain Analysis

Our FNC matrix cross-domain analysis provides a more detailed view of the link between structural and functional data. The produced and real FNC matrices have a total similarity measure of 0.74, which shows that there is a significant relationship, but the structural data does not fully reflect all functional features. The complicated nature of brain functionality, which cannot be entirely extrapolated from structural imaging, may account for this disparity.

Upon examining the cross-domain correlations, it becomes apparent that the within-domain correlations, such as AUD-AUD, exhibit a remarkably high similarity (0.955). This suggests that the structural data accurately reflects the functional connection of the auditory network. This is supported by strong correlations in subdomains like SC-AUD (0.847) and SC-SM (0.824), which show stable structural-functional alignment in the motor function and sensory processing domains.

The cViT-GAN model captures the cerebellum’s constant functional patterning, which is frequently underrepresented in FNC research, as evidenced by its strong intra-domain correlation (CB-CB at 0.821). Cross-domain interactions, like those between the default mode network and the cerebellum (DMN-CB) and the somatomotor and cerebellar regions (SM-CB), have moderate to high correlations. This means that the model can show how different parts of the brain work together.

Notably, the lower correlations in coupling between other regions including SC-CB (0.160) and AUD-CB (0.053) show the challenge of mapping functional networks from structural data, especially when there are complicated connections between regions. These areas may indicate distinct functional characteristics or dynamic interconnections that are not readily apparent in structural MRI data. These correlations are specific across domain sizes, from the small 2×2 matrices to the large 17×17 matrices. This makes them useful for checking the authenticity of FNC representations that have been made. Furthermore, it identifies areas where the generative model’s performance could be enhanced to more accurately replicate the intricate tapestry of human brain connectivity.

Finally, our findings highlight the benefits and drawbacks of employing cEViT-GAN to replicate FNC matrices using structural data. The model’s high fidelity in some domains encourages its use in clinical settings, whereas inequalities in others call for further research into the multidimensional nature of brain structure-function interactions.

### D. Structural-to-functional Connectivity

The identification of biomarkers in SZ by combining structural and functional neuroimaging data tells a captivating story about the disorder’s neuropathology. The findings of our investigation show a significant agreement between structural and functional indicators, highlighting the complicated connection between brain structure and function in SZ.

Significant structural sections include the medial frontal cortex (mPFC), dorsolateral prefrontal cortex (DL-PFC), and cerebellum. Similar functional regions show significant changes in connection patterns, particularly in the default mode network (DMN) and auditory and somatomotor activities. This correspondence between structural changes and functional connectivity disturbances allows for a more comprehensive understanding of SZ pathophysiology. For example, functional connectivity disruptions in the mPFC and DL-PFC, which are important for executive functioning and cognitive control, coincide with structural alterations in these areas, contributing to the cognitive and affective dysregulation seen in SZ patients. Both structural and functional findings highlight the importance of the cerebellum in SZ, an area that has been understudied until now. Changes in cerebellar areas correspond structurally with changes in functional connectivity within the cerebellum and its linkages to other brain networks. This shows that the cerebellum may play a role in the larger network dysfunctions that characterize SZ, going beyond its traditional concept of motor control.

Furthermore, the temporal lobe, a region involved in auditory processing, exhibits both structural and functional abnormalities, which correspond to clinical symptoms such as auditory hallucinations, which are common in SZ. This is supported by the significant correlation in functional networks, including the auditory cortex (AUD), which mirrors the anatomical findings. These similarities hint at a more integrated model of SZ in which structural anomalies are not isolated but have a considerable impact on the functional network dynamics. This model supports the idea that SZ is a disorder of “disconnected connection,” with the symptoms being caused by the interaction of damage to the structure and problems with the way the network works.

In conclusion, the convergence of structural and functional biomarkers in our work have provided some new insights into our understanding of SZ. It demonstrates the interrelated nature of structural and functional network changes, providing a more comprehensive view of the disorder’s neurobiological roots. We hope our understanding can be further increase by an integrative approach like this, pontentially leading to the development of more effective diagnostic tools and targeted treatment options that are tailored to the personalized nature of SZ.

## VI. Discussion and Conclusion

We used sMRI data to build comparable FNC matrices in our study, revealing the link between structure and function. Notably, we identified critical locations in sMRI using ViT attention maps, resulting in produced FNCs that closely mirror real data. between addition, our work advances our understanding of the structural-functional links between schizophrenia (SZ) and healthy controls (HC). The cEViT-GAN model excels in simulating actual FNC data, particularly in subcortical regions like the cerebellum and auditory networks. This not only verifies the efficiency of our approach, but also emphasizes the multifaceted character of SZ.

However, when addressing the group difference FNCs (SZ-HC), an area of great interest emerges. In this case, our produced FNCs were less comparable to genuine FNCs than whole-averaged comparisons. Future study should focus on the reduced similarity in group difference FNCs. This feature emphasizes the complicated dynamics of brain function changes in SZ patients and HC. It implies that, while our model is capable of capturing individual FNC patterns, capturing the nuanced distinctions in group dynamics is a more difficult task. This finding begs for additional exploration and model development in order to improve its value in differential diagnoses.

Despite this constraint, our research makes a substantial contribution to the discipline. It provides the path for future research to investigate the use of produced FNC matrices in classification tasks, potentially improving SZ diagnosis precision. The incorporation of our model into diagnostic procedures may result in more accurate detection of SZ, allowing for the creation of tailored therapy methods and, ultimately, better patient care in mental health.

In conclusion, our work not only increases our understanding of SZ but also offers up new research options. With its potential to expose intricate brain network patterns, the cEViT-GAN model is a significant tool in the search to understand the complexities of SZ. This paradigm has enormous potential to revolutionize the landscape of SZ diagnosis and treatment, providing a light of hope for future advances in mental health care.

## Notes

### Competing Interest Statement

The authors have declared no competing interest.

